# Integrated genetic and metabolic characterisation of diverse Latin American cassava (*Manihot esculenta* Crantz) germplasm; implications for future breeding strategies

**DOI:** 10.1101/2022.12.01.518686

**Authors:** L Perez-Fons, TM Ovalle, M Drapal, MA Ospina, A Bohorquez-Chaux, LA Becerra Lopez-Lavalle, PD Fraser

## Abstract

Cassava is an important staple crop for food security in Africa and South America. The present study describes an integrated genomic and metabolomic approach to the characterisation of Latin American cassava germplasm. Classification based on genotyping and the leaf metabolome correlates, the key finding being the adaption to specific eco-geographical environments. In contrast the root metabolome does not relate to the genotypic clustering, suggesting different spatial regulation of this tissue’s metabolome. The data has been used to generate pan-metabolomes for specific tissues and the inclusion of phenotypic data has enabled the identification of metabolic sectors underlying traits of interest. For example, tolerance to whiteflies was not linked to cyanide content but to cell wall related phenylpropanoids or apocarotenoids. Collectively, these data advance the communities resources and provides a valuable insight into new parental breeding materials with traits of interest directly related to combating food security.

**Significance statement:** Cassava is a staple crop in developing countries of sub-tropical regions. Traditionally, cassava has been considered as a subsistence crop. However recently it has become a sustainable solution to fulfil both hunger and malnutrition needs, and drive economic development. Varietal improvement via classic breeding has successfully delivered products into the Asian market by including/exchanging germplasm from original Latin American collections. Conversely, modest progress has been achieved in Sub-Saharan countries since genetic resources are biased towards exploitation of local landraces and uncharacterised parental material. The present work explores the genetic and metabolic diversity of Latin American cassava’s genebank, one of the largest and most complete worldwide. These data provide a robust characterisation of valuable germplasm that can be exploited in breeding programmes.

## Introduction

Cassava (*Manihot esculenta Crantz*) is an important staple food crop for over 800 million people in Africa and South America (1). It has also been recently proposed as a solution to circumvent the global cereal shortages arising (https://www.theguardian.com/global-development/commentisfree/2022/may/12/cassava-nigeria-wean-world-off-wheat). Contrastingly in Asia the demand for cassava has changed from a direct food crop to an industrial feedstock being processed into animal feed and starch (2). In comparison to other crops cassava is resilient to environmental fluctuations, it will grow on poor soils and agronomic production does not require sophisticated technology. Thus, it is a good target crop for addressing food and nutritional security concerns in the face of changing climates (3).

Over the last decade investments in cassava as a food system have resulted in improved tools and resources for the breeding of new varieties. This has led to the development and deployment of cassava varieties with improved yields (2) and nutritional content (4), as well as disease and pest resistance (5). However, despite these notable advances production levels will not be sufficient to impact upon the predicted global food and nutritional issues (6). In addition, it has become evident that new varieties with improved agronomic traits have experienced inconsistent adaption rates due to altered end-user preferences for various quality traits (7).

Although modern genomic selection approaches have been incorporated into cassava breeding pipelines, the fundamental strategy still mainly relies on recurrent phenotypic selection, as populations are advanced. Virtually all cassava cultivation uses clonal propagation and in most cases selfing of uncharacterised material. Collectively, this has been shown to cause inbreeding depression. For example, it has been determined that through a single inbreeding generation over a 60% decrease in fresh root yield can occur (8-10). Similar scenarios have arisen with potato breeding where cultivars are propagated using seed tubers, making the crop recalcitrant to the use of molecular/genomic approaches (11). The potato industry is now adopting a move from clonally propagated tetraploids to true seed-propagated diploids. Sexually propagated species have reduced inbreeding depression, because of deleterious mutations are not readily inherited compared to clonal propagation (12).

Having well characterised germplasm collections of cassava that are true to type, offer the potential to select suitable parental material for the construction of pre-breeding populations, capable of delivering diverse quantitative agronomic and quality traits that can be readily selected. Immortalising these lines also offer the potential for true seed.

Numerous studies exist that have used genomic approaches to genotype existing diversity. Typically this approach is restricted to local landraces collections or siblings of bi-parental populations from uncharacterised material (13, 14). However, collaborative efforts between Asian and Latin American genebank collections has resulted in successful breeding outputs and deployments of stable varieties with consistent phenotypes (2). Recently, this strategy has been progressively incorporated into pre-breeding programs to deliver biotic resistance in Africa (15-17). Among the global cassava germplasm collections available CIAT offers the largest and most diverse (1, 18). In the present study, genotypic and phenotypic data on the genetic architecture and agro-ecological profiles of the populations has been integrated with metabolome analysis. The metabolome becoming the biochemical output of the genome and often directly associated with the phenotypic traits.

Collectively the data has contributed to better characterised genetic resources, which can be used to select improved parental materials for the construction of pre-breeding populations. The outputs have also shed light on the spatial regulation of the cassava metabolome, the influence of agro-ecological adaption on the metabolome and finally identified potential targets for New Plant Breeding techniques to rapidly incorporate traits of interest into suitable cassava metabotypes.

## Results

### 1. Genetic diversity and population structure of LA cassava germplasm collection

Cassava’s genebank held at CIAT station covers a wide diversity of accessions (Suppl. Fig. 1) collected from a range of Latin America locations and biomes. A subset of 481 accessions with complete passport data (Suppl. Dataset 1) was selected for studying the genetic diversity. Bayesian analysis implemented with fastSTRUCTURE was run on the dataset without any prior classification to unravel genetic composition, genotype relatedness and population structure. The clustering method differentiate two main gene pools grouping 7 genetic sub clusters (Fig. 1A). The genetic subgroups defined as dry and humid Atlantic Forest constitute the cassava’s gene pool of south and south-eastern area of the Amazon River basin, and the gene pool corresponding to the north and north-western areas of the Amazon River includes the genetic subgroups defined as Andean high and lowlands (AHL and ALL), Amazon River basin (ARB), savanna (SAV) and Meso America (MAM). Despite the clear division between South/North Amazon River gene pools, heterogeneous presence of the different genetic subgroups is observed in the Central America and Caribbean regions (Fig. 1E). Additionally, genetic relatedness based on genetic distance indicates that MAM, SAV and AHL subgroups are closer to ARB, and the ALL partially clusters with southern subpopulations of the dry and humid Atlantic forest (DAF and HAF) (Fig. 1B). Noteworthy, a large proportion of the collection remained unclassified due to either unavailable or low quality of sequencing data (Suppl. Dataset S1).

**Figure 1:**
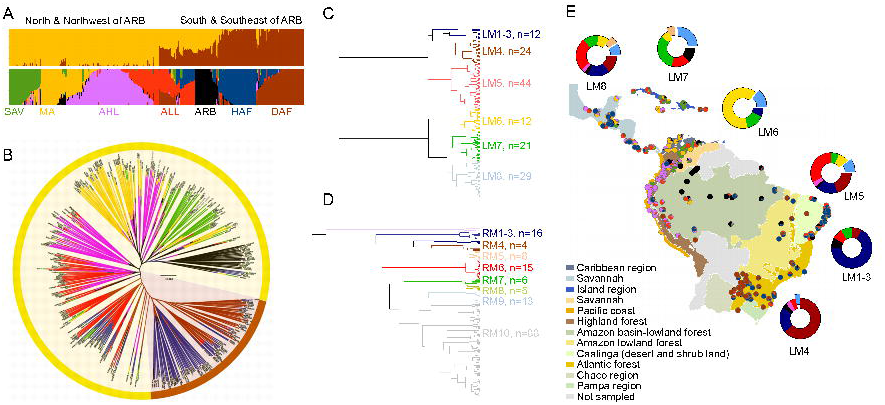
Genetic and metabolic diversity. Panel **A** illustrates the proportion of the genome of each Latin American cassava landrace assigned to each of the seven sub clusters (bottom) grouped under the main genetic gene pools (top); and (**B**) plots the group assignment upon the results of the STRUCTURE analysis. Each individual accession is represented by a vertical bar showing the genomic proportion from each sub cluster: savanna (SAV, green), Meso America (MA, yellow), Andean highlands (AHL, pink), Andean lowlands (ALL, red), Amazon River basin (ARB, black) humid Atlantic Forest (HAF, blue) and dry Atlantic Forest (DAF, brown). Dendrogram of leaf (**C**) and root (**D**) metabotype classification obtained by hierarchical cluster analysis of LC-MS untargeted data. Clusters of accessions presenting similar metabolite fingerprint are designed as leaf metabotype (LM) or root metabotype (RM) clades followed by the number of accessions grouped. Genetic subgroup’s composition of LMs are represented as doughnut plots and average proportions of the seven cassava genetic subpopulations are mapped as filled-in pie-plots over the Central and South America biomes indicated in the figure legend (**E**). LMs sectors corresponding to Asian and African accessions are coloured as pale blue and brown, respectively.

Both gene pools and genetic subgroups co-localise with the different eco-geographic regions characteristic of Central and South America (Fig. 1E). The HAF subgroup includes essentially Brazilian lines with samples from Colombia and Central America (Guatemala, Mexico, and United States), whilst the composition of the DAF cluster is shared between Brazilian, Paraguayan, Argentinian and Bolivian landraces. The ARB genetic subgroup comprise accessions from Brazil and Colombia, and the AHL cluster contains varieties from Ecuador, Peru and Colombia mostly, but also some collected from Bolivia, Mexico and Caribbean region. The largest proportion of the ALL accessions are from Brazil, Colombia and Peru, complemented with lines sampled from Ecuador, the Caribbean Islands, Central America or United States. The SAV subgroup is dominated by Venezuelan, Colombian and Cuban accessions and minor contributions from Mexico, Panama, Honduras or Ecuador. Varieties sampled from Colombia, Guatemala, Mexico, Panama and Costa Rica are the major components of the MAM sub cluster which also contained some lines from Puerto Rico, Jamaica and Venezuela.

### 2. Metabolic diversity

In the present study, an untargeted metabolomics approach has been used in order to capture the plant’s chemical diversity by including all chemical features detected without prior knowledge of their identification.

The biochemical diversity of cassava was evaluated in both uncooked roots and leaves of a germplasm sub-collection that included accessions collected from Central and South America and a limited representation of African and Asian varieties. A number of advanced lines developed under CIAT’s breeding projects was also part of the diversity panel screened.

Classification of accessions based on chemical fingerprint similarity was generated by hierarchical clustering analysis using untargeted metabolomics data as input matrix (Suppl. Dataset S2, S3). The resulting dendrograms differentiate 8 clusters of accessions in the leaf tissue (Fig. 1C) and 10 clusters of accessions in root tissue (Fig. 1D). The groups of accessions were nominated as leaf-metabotype (LM) or root-metabotype (RM) clades, respectively. The number of cassava accessions were homogenously distributed along the different LM clades, whilst the number of samples in every RM clade was unequally distributed with RM-10 being the largest and concentrating approximately 55% of the accessions sampled. In addition, inconsistency between leaf-metabotype and root-metabotype classification is evident, and therefore the implications in relation to genetic diversity and phenotype are subsequently analysed and discussed separately.

#### (i) Leaf metabolome diversity

The classification of leaf-metabotype clades mirrors the diversity of genetic groups extracted from the SNPs analysis and their corresponding eco-geographic locations (Fig. 1E). Overall, the LMs 1 to 5 comprise accessions of the South and South-east Amazon River Basin gene pool, whilst the bottom branch of the LM-dendrogram concentrate those accessions genotyped as North and North-western Amazon River Basin gene pool. Leaf metabotypes 1 to 4 group cassava accessions classified under the HAF and DAF genetic subgroups, mapped in the south and south-east areas of the Amazon River towards the Atlantic coast. In addition, LM5 presents almost equal contributions of ALL accessions and Atlantic Forest accessions collected from these southern regions. Approximately three quarters of LM6 accessions fall within the MAM genetic sub-cluster, with Colombian lines from the areas facing the Caribbean Sea being the largest contributors. Landraces collected from Venezuela, Cuba and Panama form the savanna representatives of LM7, and Brazilian samples contribute to the Amazon sector of this clade. Finally, LM8 shows a similar composition of genetic backgrounds as the southeast cluster LM5, hence nominated as Mixed-North and Mixed-South clades, respectively. The difference between both mixed LM clades being the geographical locations from where the samples were collected. Leaf clade LM5 is dominated by Brazilian and Colombian landraces, whilst LM8 includes lines from Venezuela, Guatemala, Peru and Paraguay. Noteworthy that African accessions cluster under LM7-8, and Asian samples spread over LMs5-8 and predominantly in LM7, although genetic classification was not available for these foreign accessions.

The number of chemical features significantly differentiating each LM clade is also annotated in the metabotyping dendrogram (Suppl. Fig. 2). The two main South/North branches differ in 403 chemical features, and 50 and 87 mass signals differentiate the sub-clades within South and North respectively. Savanna and Meso America accessions (LM7 and 6), separate out by 72 differentiating chemical features, and 290 metabolite signals are significantly different between leaf mixed-clades LM5 and 8, despite both presenting similar composition of genetic backgrounds.

**Figure 2:**
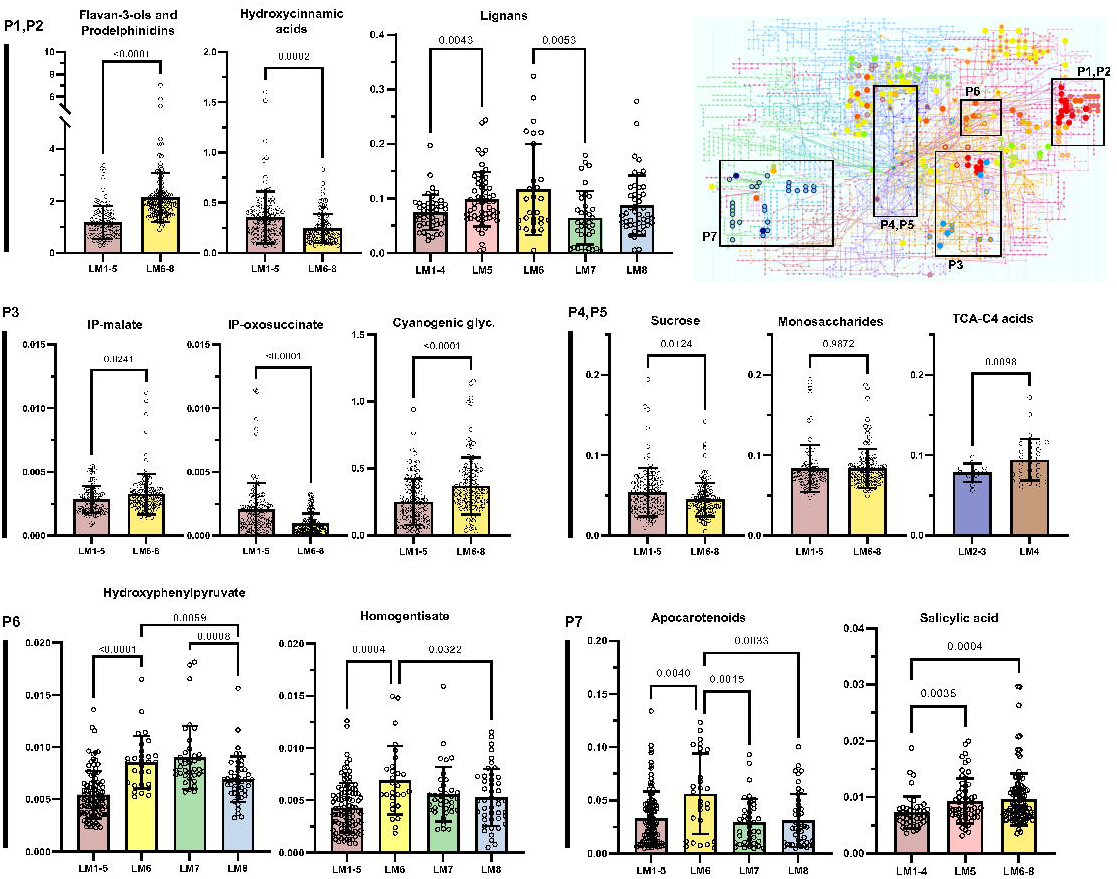
Enriched metabolic pathways differentiating cassava’s LM clades mapped onto KEGG’s *Arabidopsis thaliana* metabolic network (top-right). Examples of metabolite’s relative concentrations (to internal standard) of some of the most significant enriched pathways are displayed in single bars plots representing mean, standard deviation and individual values. Pair-wise p-value is indicated in each graph. Level of statistical significance between LM clades was obtained from one-way ANOVA analysis (unpaired with Welch’s corrections and Brown-Forsythe test for unequal variance) and multiple pair-wise comparisons adjusted with a family-wise threshold (alpha) of 0.05 (95% confidence interval). P1, P2: Hydroxycinnamic acids, flavonoids, and lignan pathways; P3: Valine, leucine and isoleucine metabolism leading to cyanogenic glycosides biosynthesis; P4, P5: energy metabolism including sugar metabolism and TCA cycle. C4-acids included malic and fumaric acids. P6: Tyrosine metabolism and quinone ring biosynthesis; P7: phytohormones. Colouring of clades follows legend of LM dendrogram as displayed in figure 1. LM: leaf metabotype clade.

#### (ii) Root metabolome diversity

The metabotyping classification tree based on the analysis of root tissue provided with 10 clades of root metabotypes (Fig. 1D). The RM clades are arranged in two main clusters, the top one grouping clades 1 to 4 of which over 50% of accessions include humid and dry Atlantic Forest representatives. However, inconsistent alignment of root-metabotype (RM) clades and genetic subgroups, LMs or eco-geographical biomes is observed (Suppl. Fig. 3). The exceptions are RM6 and 7 of which composition of genetic groups concentrate over 75% of Andean lowlands and savanna accessions, respectively. Nevertheless, the total number of accessions included in RM6 or RM7 represents less than 10% of the collection. Similarly, RM9 shows similar genetic groups composition as RM1-4 with additional contributors from the ARB sub-cluster, but no consistency in geographical locations sampled mark the difference between these two distant RM lades.

**Figure 3:**
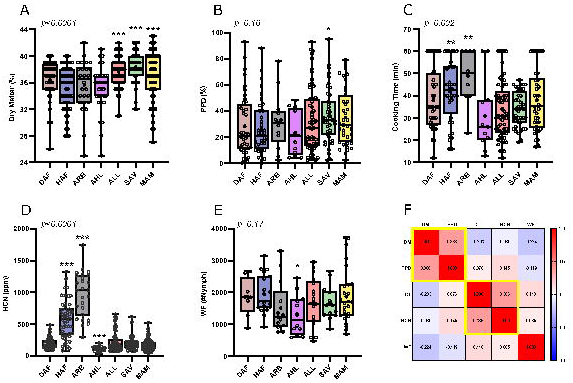
Box plots of phenotyping data collected from 2014 to 2018 at CIAT station. Bold central line and filled-in circle indicate median and average values respectively, and accessions’ individual values as open circles. **A**, DM: dry matter in %; **B**, PPD: % of post-harvest physiological deterioration of root; **C**, cooking time of roots in minutes; **D**, HCN: roots’ cyanide content in parts per million (ppm); **E**, WF: leaf nymph counts of whitefly *Aleurotrachelus socialis*; and **F**, matrix Pearson’s correlation between phenotypes measured. Yellow boxes denote significant (p<0.05) correlations. DAF: dry Atlantic Forest (brown); HAF: humid Atlantic Forest (blue); AHL: Andean highlands (pink); ALL: Andean lowlands (red); SAV: savanna (green); MAM: Meso America (yellow). Phenotype’s ANOVA (one-way) p value is indicated at the top of every plotting area, and statistical significance of pair-wise comparisons between genetic sub groups is also labelled over the corresponding significant group: p<0.001(***), p<0.01 (**), p<0.05(*).

### 3. The construction of a pan-metabolome for cassava

In line with the pan-genome definition, “ the entire set of genes from all strains within a clade”, a homologous description is proposed to the pan-metabolome term as the entire set of metabolites present in all natural variants within a clade. Thus, the unbiased nature of untargeted metabolomics together with the wide variety of natural accessions analysed has enabled the characterisation of cassava’s pan-metabolome in the present study. Unlike genomic (DNA) information, metabolomic information is tissue specific and therefore a spatial metabolomics approach has been followed by analysing both aerial and storage tissue (leaf and root). The capture of the entire metabolome is feasible thanks to the operational essence of TOF (time-of-flight) mass analysers which facilitates the detection of all chemical components present in the matrix analysed. As opposed to the studies based on triple quadrupole technologies which rely on targeted detection of known compounds pre-selected by user.

The main bottleneck of untargeted metabolomics data analysis pipelines is the identification and annotation of the chemical entities detected. Here, two alternative workflows were followed for metabolite identity characterisation of the leaf and root pan-metabolomes. Mining of the leaf metabolome was performed by combining pathway enrichment analysis of significant differentiating features and annotation of untargeted matrix using in-house metabolite libraries validated with chromatographic and mass spectral parameters. This will allow elucidation of diversity at genetic and metabolic level in leaf tissue since both LM-clades and genetic groups’ classification mirror each other. On the other hand, root metabolome elucidation was arranged by annotating the top most abundant molecular features characteristic of every RM clade using accurate mass measurements.

#### (i) Pan-metabolome of leaf tissue

Pathway enrichment analysis was performed by using the outputs of pair-wise comparison between the different LM clades or groups of them as input data (Suppl. Dataset S4). Results of these analysis are summarised in Suppl. Table 1 indicating the most significant enriched pathways, and full detail of all pathways and metabolites enriched per pair-wise comparative are included in Suppl. Dataset S5. Highlights of the results are visualised in Fig. 2. Accessions from the Southern (LM1-5) and the Northern (LM6-8) Amazon River regions differ in the composition of hydroxycinnamates, flavan-3-ols and their polymeric forms (prodelphinidins), but no significant differences were observed in the flavone and flavonol composition. Condensed tannins prodelphinidins were higher in the Northern LM-clades and opposite trend was found for their intermediaries, the hydroxycinnamates. Subtle significance was observed in the sucrose levels between the North *versus* South LM-groups, but stronger significant differences were detected in components of the aliphatic amino acids metabolism valine, leucine and isoleucine which ultimately feed into the cyanogenic glycosides biosynthesis of linamarin and lotaustralin. Comparatives within LM clades forming South and North dominant groups retrieved distinctive sections of the metabolome too. For example, significant changes in the TCA cycle-related organic acids were found between representatives of HAF and DAF (LM1-3 *vs*. LM4); or different levels of lignans were significant between South-mix clade LM5 and LM1-4 accessions. Similarly, the lignan content of LM6 was the highest over the rest of LM clades. The clade LM6, largely representing MAM accessions, also presented elevated relative concentrations of apocarotenoids and components of the tyrosine metabolism, 4-hydroxyphenylpyruvate (4-HPP) and homogentisate, the latter leading to the biosynthesis of tocopherol’s chromanol ring. Finally, salicylic acid and its glycosylated form were significantly more abundant in the Northern LM-clades (LM6-8) and the south-mix LM5 compared to the rest of Southern clades LM1-4.

#### (ii) Pan-metabolome of root tissue

The complexity of the chemical composition of root extracts increases from the accessions located at the center of the PCA score plot towards extremes (Suppl. Fig. 4A). A large group of accessions included in RM10 (grey) were characterised by lower chemical diversity, whereas clades RM1 to 9 present specific chemotype profiles with both qualitative and quantitative features. The relative quantities of chemical features and putative annotation of the ten most abundant chemical signals per RM clade are visualised in Suppl. Fig. 4B. Root-metabotype clades 1-4 and 5 show the components with the highest accumulation levels. These have been putatively characterised as lignin-type in clade 1 and caffeoyl-glycosides, hydroxycinnamate and hydroxybenzoate derivatives in RM2 to 5 (Suppl. Dataset S6). In addition, RM4 presented higher relative quantities of nitrogenated compounds and glycosyl-chalcone and methoxy-coumaroyl derivatives. In contrast, RM6 to 8 contain elevated levels of different monoterpene iridoids and other terpenoid structures in glycosylated form. Root metabotype 6 also show higher abundance of certain components involved in the urea cycle, acetyl-glutamic acid and itaconic/citraconic acids. Additional components found in RM8 were putatively characterised as glycosylated variants of gallotannins and sinapic acid. The metabotype cluster with lowest chemical complexity is represented by RM9 with only one tetrapeptide characterised.

**Figure 4:**
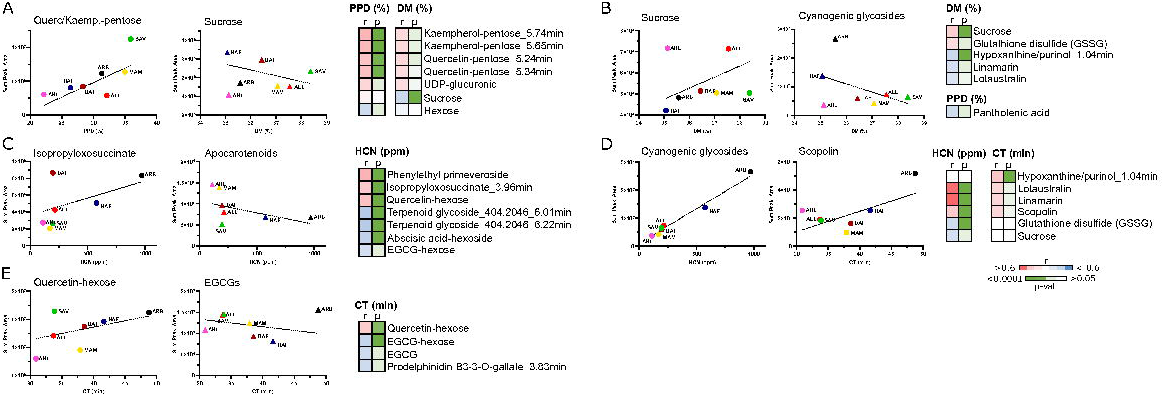
Correlation (Pearson) plots between phenotypes and leaf metabolites (**A**) or root metabolites (**B**). Accessions’ correlation (r) and significance (p) values are displayed on the left-side and average values per genetic sub-clusters on the right-side. Ppm: parts per million; EGCG: epigallocatechin gallate; DAF: dry Atlantic Forest (brown); HAF: humid Atlantic Forest (blue); AHL: Andean highlands (pink); ALL: Andean lowlands (red); SAV: savanna (green); MAM: Meso America (yellow).

### 4. Phenotype-metabolite link per genetic groups

#### (i) Quality traits and metabolome stability

Phenotypic data concerning dry matter, cooking time, physiological post-harvest deterioration (PPD) or root cyanide (HCN) content were recorded between 2014 and 2018 and metabolome stability of tubers was evaluated on material collected from 2016 to 2020 crop years. In addition, a small subset of 110 accessions was tested for whitefly (*Aleurotrachelus socialis*) tolerance as part of CIAT’s research program and included in the present study. Genetic subgroups differ significantly in dry matter, cooking time and cyanide content (Fig.3A, C, D), whilst no significant differences were detected for PPD or whitefly tolerance (Fig. 3B, E), the latter likely due to the size of the subset being insufficient for assessing statistical significance. Nonetheless, subtle significant differences were found in the pair-wise comparisons between genetic groups with AHL being the most tolerant to whitefly (lowest count) and the lowest in root HCN. Amazon and humid Atlantic Forest groups presented the highest values of cyanide content and longest cooking times, whilst Andean lowlands, savanna and Meso America genetic subgroups show the highest levels of dry matter. Correlation matrix of phenotypic data paired per accessions indicates modest but significant correlation between DM and PPD or HCN and CT (Fig. 3F).

The principal component analysis (PCA) score plot including all accessions and crop years explained 10% of variation and shows a consistent overlap between crop years (Suppl. Fig. 5A). In order to assess metabolite composition stability over the years, a subset of 189 accessions present in both 2016 and 2020 crops was compared using orthogonal partial least square discriminant analysis (OPLS-DA) (Suppl. Fig. 5B). The OPLS-DA model presented modest fitting (R2) and cross-validation (Q2) values of 0.609 and 0.469 respectively. Variation between classes (years) was 0.0233 and within class (genetic diversity) was 0.0443 (Suppl. Fig. 5B).

#### (ii) Phenotype-metabolite link

Correlations between accessions’ phenotypic records and leaf and root’s chemical features were attempted as proof-of-concept application of the genetic and metabolome resources generated in the present study.

Pentose derivatives of leaf flavonols quercetin and kaempherol significantly correlate with PPD and DM phenotypes in a positive manner (Fig. 4A), and whilst negative correlation is observed between leaf-sucrose and hexose, positive correlation is observed between these quality traits and root-sucrose (Fig. 4B). Similarly, levels of linamarin and lotaustralin in roots positively correlate with HCN and CT traits but not with leaf levels of the same cyanogenic glycosides (Fig. 4A, B). In addition, the flavonol quercetin-hexose and the flavan-3-ol epigallocatechin gallate (EGCG) and its derivatives (hexoside and polymeric forms) in the leaf negatively correlate with both HCN and CT. On the other hand, the coumarin scopolin did not correlate with PPD but with HCN and CT phenotypes in root’s tissue, and only pantothenic acid negatively correlated with PPD (Fig. 4B).

Additionally, a number of uncharacterised chemical features in both leaf and root tissue show strong and significant correlation with the different phenotypic traits studied (Suppl. Datasets S7, S8).

The correlation approach was utilised to identify potential metabolite markers of whitefly tolerance in the cassava’s Latin American collection. No significant correlations were obtained from the analysis of the whitefly phenotype and leaf metabolite chemical classes, except for apocarotenoids. This group of compounds, including ABA and its derivatives, and putative blumenols, positively correlate with whitefly nymph count and negatively with prodelphinidins and lignans, although below statistical significance (Suppl. Fig. 6). Overall, these results suggest that whitefly tolerance is a complex trait involving multiple metabolome sectors likely modulated by signalling molecules, and therefore detailed characterisation of the biochemical and molecular mechanisms of action would be required to fully elucidate potential phenotype markers.

## Discussion

Although large-scale genomic studies on cassava diversity exist, the present approach represents one of the largest metabolomic study performed on cassava to date. Untargeted LC-HRMS/MS was deployed because of its unbiased nature, mass accuracy, MS/MS capability and ability to accurately capture wide diversity of chemical features. Thus, in comparison to other analytical apparatus it provides the closest reflection of the metabolome that can be achieved using one analytical platform. The overriding feature of the genotyping data across the CIAT diversity panel was the clustering on agro-ecological location, previously reported in smaller scale studies (19-22). In the case of the leaf metabolome there was a clear correlation with the genotypic data. This suggests a direct flow of information from gene to metabolite supporting the central dogma of biology. However, the analysis of diversity and stability of the root metabolome indicates that the genetic regulation of this tissue overrides the effect of adaptation pressure and environmental conditions.

The integrative analysis presented in this study also suggests to the presence of two centres of domestication for cultivated cassava in Latin America; maybe evolving from different ancestors and coincidental with regions of intensified breeding activity, Central America, including Caribbean regions, and South-eastern Brazil. Whilst introgression regions from wild *Manihot glaziovii* are evident in Brazilian and African germplasms, the original ancestor of Colombian and Northern regions of Amazon basin remain unclear and are still under debate, but presumably from *Manihot flabellifolia* (21, 23). Hence the presence of two distinctive gene pools within the Latin American collection, each comprising independent metabolome clades of mixed genotypes. It is interesting that cassava diversity evolution, follows a similar pattern to the Solanaceae crops such as tomato or pepper (24, 25). The data also highlighted the importance of genetic and biochemical adaption to the environments present in these specific agri-geographical locations. Presumably this phenomenon is one of the reasons why germplasm of Latin America (LA) origin cannot readily be cultivated in other biomes (19), such as those found in Sub-Saharan Africa. To overcome these limitations but utilise the LA diversity for traits of interest, genetic crossing into an intermediate background could offer potential as precedents exist (2). For example, some Latin American accessions classified under the domesticated clade LM8 (e.g., VEN164, VEN173) have been reported genetically close to the Central East African germplasm; or the LA accession CG1320-10 (a cross between MEX1 and PAN51) which is widespread in Africa and genetically clusters under the African breeding germplasm (18). It could also be postulated that the genotypes from diverse agro-ecological backgrounds offer potential as genetic donors for biotic and abiotic stress resistance predicted to arise from climate change parameters (19).

Interestingly classification of the root metabolomes did not reflect the genotypic classification of the leaf metabolomes. This suggests that different spatial and temporal regulation of the root metabolome in cassava occurs, reflecting its biochemical specialism. In the case of cassava, the tuber is the edible tissue of the plant and thus directly exposed to consumer preferences, which typically relate to quality. One of the issues with present breeding programs utilising genomic parameters solely is the prevalence of agronomic trait (mostly yield) over consumers’ preference traits. Typically, this is due to poor quality traits and frustratingly arises after time-consuming and expensive breeding cycles. Such findings suggest a larger metabolomic and/or sensory evaluation would be beneficial to breeding programmes. Recently, it has been shown that metabolome selection is an accurate predictor of fruit flavour in tomato complementing genomic selection and sensory traits (26). Based on these data the inclusion of metabolomic selection into the breeding programme is proposed. In the case of cassava, the tuber metabolome does not correlate with the genotypic classification used in this study but correlation with consumer and agronomic traits has been stablished (27). Therefore, the inclusion of metabolome analysis and sensory evaluation is necessary if new varieties with acceptable quality traits are to be produced.

The spatial differences in the leaf and root metabolomes across this diverse panel also imply the population represents a good resource for the study of source-sink genetic elements. The data generated represents an advance to the genetic/biochemical resources available in cassava. For example, the present study has facilitated the further identification of metabolites by clustering of genotypes, and enables future association studies using GWAS approaches (28-30). Similar approach has been applied in tomato and maize (31, 32). In addition, the study has generated a version of the pan-metabolome for cassava, enabling a core collection of metabolites in cassava which reduces or focuses the number of key molecular features requiring intense identification.

The incorporation of selected traits into the dataset has enabled the identification of specific clades showing enhanced quality or agronomic traits. The addition of the metabolomics data facilitates deciphering the underlying biochemical and molecular mechanism associated with the traits of interest and how this is linked to environmental adaption. One of the examples used is the resistance (or tolerance) of cassava to whitefly infestation. Traditionally, it is accepted that cyanide containing cassava had evolved as a strategy to alleviate biotic stresses. However, in the case of whitefly infestations the leaf cyanide content does not correlate with the lowest count of whitefly but with signalling molecules modulating cell wall-related phenolics. The latter corroborating the mechanism postulated previously (15).

In summary the present data advances the resources in cassava and is an example to other clonal propagated crosses. The accessions characterised can act as parental material in future breeding activities, where the true to type nature can pave the way for the long-term goal of true seed that will reduce future breeding depression by facilitating cultivation from seed stocks. Although there is still much to learn and discover, this study shows the benefits of applying metabolomics to genotypic capture of diversity and potentially subsequent genome assisted plant breeding. Conversely, it also demonstrates how genomics can have an important impact on expanding our knowledge of biochemical pathways and underlying mechanisms associated with traits. Such mechanistic elucidation will aid more rationale design of breeding approaches in the future.

## Methods

### Plant material and growing conditions

Based on the geographical distribution of 3,331 LAC landraces held at CIAT cassava world collection, we randomly selected an experimental population of 481 individuals to account for the most heterogeneous cassava landraces. These cassava landraces distributed throughout North, Central and South America were transferred from tissue culture into a screen-house for tissue hardening (33). All agronomic data was collected at CIAT experimental extension with an elevation of an approximately 900 m above sea level. This population was screened for agronomic performance across four growing seasons (2014–2018). The growing season typically started in May and harvest the following year in March/April. These trials were designed following a random complete block design (RCBD) with three replications (plots); each plot consisted of a total of 16 plants. The agronomic data were collected plant-by-plant, for the four inner most plants, in each plot 10 months after planting (MAP). The plots with less than 10 geminated plants were removed before data analysis.

### DNA extraction and Next Generation Sequencing (Whole and RAD-Seq)

DNA was extracted according to the CTAB-based DNA extraction protocol described by (34) with minor modifications. The disrupted tissues were incubated at 65°C for 1 h follow by one organic extraction using chloroform-isoamyl alcohol (24:1); mixing gently but thoroughly for 30 min at 0°C. The DNA resulting from this extraction was assessed for quality in a 1% agarose gel and quantified using Synergy™ HT Multi-detection microplate reader (BioTek®, USA). RAD-tag libraries were developed by the Beijing Genomic Institute (BGI) following the method described by (35) using the *EcoR*I restriction enzyme (recognition site: 5’-G/AATTC-3’). The RAD-Seq products from the 355 LAC cassava landraces were processed in the next generation Illumina® sequencing platform HiSeq2000 (at BGI, Hong Kong, China).

### Genotyping

The cassava reference genome v6.1 (582.25 Mb arranged on 18 chromosomes plus 2,001 scaffolds) was downloaded from the Phytozome website (www.phytozome.net), including the corresponding GFF3 files containing functional gene annotations (36). The GATK pipeline was used to map RAD-Seq reads against the cassava reference genome for discovering SNPs and small InDels (37). NGSEP uses Bowtie2-2.1.0 for read alignment; except for the maximum number of alignments per read (set at 3), and the minimum and maximum fragment length for valid paired-end alignments (set between 175 to 400bp), all other parameters were run on default settings (38). For the detection of single nucleotide SNPs, NGSEP was used on default settings: 1) minimum genotype quality (40); 2) maximum value allowed for a base quality score (30); and 3) maximum number of alignments allowed starting at the same reference site (100). We first filtered SNPs in repetitive regions of the genome follow by filtering out SNPs genotyped in less than 85% of the samples; and with Phred quality scores higher than 40 (Q40) in each sample. We then excluded indels, multiallelic and monomorphic variants, leaving only biallelic SNPs variables in the final data set. Finally, we excluded SNPs with other variants within 10bp. The filters, functional annotation of variants, and the conversion of variant call format (VCF) to other formats for downstream analysis were performed also in NGSEP (37). We used STRUCTURE and PCA to analyse genetic structure patterns of the cassava accessions; both analyses were undertaken using 12,000 SNPs. The ΔK method was used to estimate the number of genetic clusters (39). The raw RAD-sequencing reads have been submitted to NCBI SRA (study accession number PRJNA245184) (http://www.cassavabase.org/).

### Phenotyping

In 2013, the 481 cassava genotypes hardened in the screen-house were transferred to the field at CIAT (Palmira, Colombia) to generate 70 stem-cuttings. These stem cuttings were used for the 2014 trial, and every subsequent year stems cutting from the year before were used for the 2015, 2016, 2017 and 2018 cassava growing seasons at CIAT’s experimental station. For each genotype, 16 stem cuttings were planted at isolation plots at a spacing of 1 m x 1 m. Each 1 m x 1m plots was separated by 1 m from any other cassava plot to ensure that plant competition was accounted for among the same genotypes. The layout of the experiment was in a complete randomized design with 4 replications under the same experimental conditions.

At harvest, which coincided with 11 months after planting, four innermost plants per genotype were uprooted and used for phenotypic assessments. Roots were separated from the vegetative harvestable biomass (leaves, stems, and original planting stake) and independently weighed. Estimation of dry matter content (DMC) and cyanogenic potential (HCN) was performed as described by (40). For the post-harvest physiological deterioration (PPD), we implemented following the methodology described by (41) and cooking-time experiments were conducted as described by (42).

### Tissue collection and preparation for metabolite analysis

Leaves and roots were separately collected and immediately frozen in liquid nitrogen. Frozen tissue was then freeze-dried for 2-3 days and ground to fine powder as described in (43). Samples were then stored at -20° C until analysis.

### Metabolites extraction

Ten mg of freeze-dried ground tissue was utilised for extraction of metabolites as described in (15). Briefly, 700 μl of 50% methanol was added and the mixture shaken for 1 hr at room temperature. Addition of 700 μl of chloroform followed by centrifugation (3 min, 14000 rpm) allowed separation of semi-polar and non-polar compounds into the epiphase and organic phase respectively. Semi-polar extract was filtered with 0.45 μm nylon membranes and non-polar extract dried under vacuum. Both extracts were kept at −20° C until analysis.

### Untargeted metabolomics analysis by LC-MS

An aliquot of 95 μl of the semi-polar extract (epiphase) was transferred to glass vials and spiked with 5 μl of internal standard (genistein at 0.2 mg/ml in methanol). Samples were kept at 8° C during analysis and volume injection was 5 or 1 μl for root and leaf extracts respectively.

For the analysis of the semi-polar extracts a C18 reverse phase column and a UHPLC-ESI-Q-TOF system from Agilent Technologies was used. The analytical platform consisted of a 1290 Infinity II liquid chromatograph, a 6560 Ion mobility Q-TOF mass spectrometer operating in Q-TOF mode only and equipped with an Agilent Jet Stream (AJS) electrospray. Data was acquired in MS mode from 100 to 1700 mDa under negative electrospray ionisation. Nebulizer and sheath gas temperatures were 325° and 275° C respectively; flowrate of drying and sheath gas (nitrogen) were 5 and 12 L/min respectively and nebulizer pressure 35 psi. Capillary VCap, nozzle and fragmentor voltages were set up at 4000, 500 and 400 V. A reference mass solution was continuously infused to ensure mass accuracy calibration. Compounds were separated in a Zorbax RRHD Eclipse Plus C18 2.1×50 mm, 1.8 μm and two different chromatographic methods were optimised for root and leaf tissue respectively. Roots’ extracts were analysed with a gradient involving (A) 0.1% formic acid in water, and (B) 0.1% formic acid in 97.5% acetonitrile. Root’s chromatographic separation proceed from 5% B held for 1 min to 30% B in 5 min followed by steep increase to 98% B in 1.5 min. After 1.5 min at 98% B, initial conditions were restored and column was re-equilibrated for 2 min. Similarly, leaf’s chromatographic method used (A) 2.5% acetonitrile in water and (B) acetonitrile as mobile phase, both solvents containing formic acid (0.03% vol.) as additive. Gradient started at 2% B for 1 min, increase to 30% B over 5 min, stay isocratic for 1 min followed by an increase to 90% B in two minutes and stay isocratic for another two minutes. Initial conditions were restored and re-equilibration lasted 3 minutes. Flowrate and column temperature of both chromatographic methods were set at 0.3 ml/min and 30° C, respectively.

### Processing of LC-MS data files: extraction of chemical features

Retention time alignment (maximum time shift +/- 0.2 min) and extraction of chemical features was performed by using Agilent’s Profinder (version 10.0) software in batch recursive mode. The following settings were selected to extract molecular features within a retention time (RT) range of 0.3 to 12 min: peak height threshold 1000 counts, RT tolerance +/- 0.15 min, mass tolerance 10 ppm, chlorine and formic acid adducts, and water neutral losses were also considered. Only molecular features (MF) with matching scores higher than 70 and present in at least 70% of each sample group (QC and samples) were included in the final data matrix. This resulted in the detection of over 500 molecular features extracted (MFE) in root and over 2500 MFE in leaf. Putative characterisation of chemical identity was inferred from accurate mass values calculated from mass-to-charge ratio (m/z) signals. Chemical formulae was generated using the following elemental constrains: C: 70; H: 140; O: 40; N: 10; S: 5; P: 3 (44), (https://pmn.plantcyc.org/CASSAVA/search-query?type=COMPOUND&formula=C), formic acid and chlorine adducts and/or multiply charged species (z=1, 2). Those chemical formula (up to 5) with the highest score (based on mass difference and isotopic pattern fitting) were selected for blasting against ChemSpider and Dictionary of Natural Products chemical databases. Additionally, an in-house library of cassava metabolites based on chromatographic parameters (retention time) and fragmentation pattern (MS/MS) (15) was used to complement and validate findings of the putative identification pipeline described above.

### Data processing and Statistical analysis

Batch correction of LC-MS untargeted data (extracted ion chromatogram peak areas) was applied using quality control samples (QC). In addition, normalisation against area of internal standard was performed. Missing values were input by using the median value of each mass reported from the extraction chemical feature pipeline, and those presenting over 75% of missing values were excluded from analysis. The resulting data matrix was then used as input for multivariate analysis (PCA, HCA and OPLS-DA) in SIMCA v17 (Sartorius AG, Germany) and univariate analysis (t-tests, ANOVA, Pearson’s correlation) in Prism v9.4.0 (GraphPad software LLC). Centering, univariate (in PCA) and pareto-scaling (in OPLS-DA) were applied for multivariate analysis. Pair-wise comparison of leaf metabotype clades were performed by multiple two-sample t-test assuming unpaired data, Gaussian distribution (parametric) and inconsistent standard deviation (Welch test). Multiple comparisons were corrected with Holm-Sidak post hoc test setting a threshold value for significance (alpha) at 0.01. The adjusted p-values obtained from the multiple t-test analysis and the corresponding m/z values of chemical features were used as input data for pathway enrichment analysis using the Functional Analysis module (MS peaks to pathways) in MetaboAnalyst v5.0 (45). Settings selected were: negative mode, 10 ppm of mass tolerance and p-value cut-off of 0.05 for the *Mummichog* algorithm. The pathway library selected was *Arabidopsis thaliana* from KEGG.

Relative amounts of metabolites validating the outputs of enrichment analysis were plotted in Prism and calculated by dividing metabolite’s peak area by internal standard’s peak area. One-way ANOVA was applied to assess level of significance assuming unpaired Welch and Brown-Forsythe tests and multiple comparisons of groups were corrected by a family-wise threshold of 0.05. Same assumptions and tests were chosen for the statistical analysis of phenotypic records.

## Funding and acknowledgements

This work was supported by the Consortium of International Agricultural Research Centers (CGIAR) Research Program on Roots, Tubers and Bananas (RTB), with additional support from the African Cassava Whitefly Project (ACWP)-Phase 2 funded by Natural Resources Institute (NRI), University of Greenwich, UK, from a grant provided by the Bill and Melinda Gates Foundation (OPP1200124). We thank Mr. Chris Gerrish for technical support, Agilent’s UK application team for assistance with metabolomics data processing and CIAT’s bioinformatics and entomology team for assistance in the design and conduction of whitefly tolerance phenotyping experiments. We also thank members of the RTB program and ACWP project for fruitful discussions.

## Authors’ contribution

LP-F and MD perform metabolomics analysis and generated corresponding dataset and related figures. TMO and AB-C perform genotyping analysis and LABL-L generated dataset and genetic classification figures. TMO, MAO, AB-C and LABL-L selected and provided plant material and conducted the phenotypic assessments. PDF and LABL-L secured funding and devised concept. LP-F LABL-L and PDF drafted the manuscript and all authors participate in discussion, interpretation of results and revision of final version. PDF and LABL-L act as corresponding authors.

## Data availability statement

All relevant data supporting results presented are provided in supplemental files. Metabolomics data sets of raw (areas) and processed data (normalised), and SNPs data are accessible in Mendeley Data and Digital Commons Data:

Perez-Fons, Laura (2022), “ EIC-Areas of chemical features extracted_Cassava diversity_LEAF tissue”, Mendeley Data, V1, doi: 10.17632/pwygzgp5rd.1. https://data.mendeley.com/datasets/pwygzgp5rd/1

Perez-Fons, Laura (2022), “ EIC-Areas of chemical features extracted in ROOT tissue of a cassava diversity collection_2016-2020 crops”, Mendeley Data, V2, doi: 10.17632/29t9bwhdmj.2. https://data.mendeley.com/datasets/29t9bwhdmj/1

Perez-Fons, Laura (2022), “ EIC-Areas of chemical features extracted_Cassava diversity_ROOT tissue_2020 crop subset”, Mendeley Data, V2, doi: 10.17632/pgwzvbm6cx.2. https://data.mendeley.com/datasets/pgwzvbm6cx/1

Perez-Fons, Laura (2022), “ QCISNormAreas of chemical features extracted in Cassava Diversity LEAF extracts_input for MVA”, Mendeley Data, V2, doi: 10.17632/s9w932ybjz.2. https://data.mendeley.com/datasets/s9w932ybjz/1

Perez-Fons, Laura (2022), “ QCISNormAreas of chemical features extracted in Cassava Diversity ROOT extracts_2016-2020 crops_input for MVA”, Mendeley Data, V2, doi: 10.17632/tgrpbpw6yz.2. https://data.mendeley.com/datasets/tgrpbpw6yz/1

Perez-Fons, Laura (2022), “ QCISNormAreas of chemical features extracted in Cassava Diversity ROOT extracts_2020 crop_input for MVA”, Mendeley Data, V2, doi: 10.17632/pbjytvmtvf.2. https://data.mendeley.com/datasets/pbjytvmtvf/1

Becerra Lopez Lavalle, Luis Augusto; Bohorquez-Chaux, Adriana (2022), “ SNPs of Cassava Diversity South American collection”, Mendeley Data, V2, doi: 10.17632/r4bn3r9k9x.2. https://data.mendeley.com/datasets/r4bn3r9k9x/1

## Figure captions

**Supplementary Figure 1:** Number of cassava accessions present in genebank collections worldwide up to 2010. Source: FAO. Save and Grow: Cassava (2014); FAO 2010. The second report on the state of the world’s plant genetic resources for food and agriculture. Rome.

**Supplementary Dataset S1 (.xlsx):** Data passport of accessions analysed and annotated according to their genetic sub group, and including phenotypic records of root dry matter, root PPD, root HCN, root cooking time and leaf nymph count of whitefly *Aleurotrachelus socialis*. N/D indicates accession of which genotype was not determined.

**Supplementary Dataset S2 (.xlsx):** List of molecular features extracted form TICs (total ion chromatograms) of leaf semi-polar extracts analysed by LC-MS. Accessions are arranged in columns and chemical features in rows including annotation of chemical identity when available. Areas of every chemical feature per sample were computed from peak integration of extracted ion chromatograms (EICs). Summary of in-house library of cassava metabolites, included as a separate sheet.

**Supplementary Dataset S3 (.xlsx):** List of molecular features extracted form TICs (total ion chromatograms) of root semi-polar extracts analysed by LC-MS. Accessions are arranged in columns and chemical features in rows. Areas of every chemical feature per sample were computed from peak integration of extracted ion chromatograms (EICs).

**Supplementary Figure 2: A**, dendrogram of leaf metabotype clades including the number of significant chemical features (DCF) differentiating each branch. Pair-wise comparison of LM-clades was performed by using multiple t-test analysis and Holm-Sidak post-hoc test (alfa=0.01). B, genetic composition of accessions clustered in every leaf metabotype (LM) clade mapped over the cassava genetic sub-groups distribution over Latin American eco-geographic regions.**Supplementary Figure 3: A**, dendrogram of root metabotype clades and pie-plots displaying the genetic composition of accessions clustered under every RM-clade. **B**, genetic composition of accessions of some RM-clades mapped over Latin American biomes when both genetic and metabotype classification match.

**Supplementary Dataset S4 (.xlsx):** Statistical analysis outputs of multiple t-test pair-wise comparison of LM clades used as input for pathway enrichment analysis.

**Supplementary Table 1 (.docx):** Summary of the most significant enriched pathways differentiating LM clades. Pathway enrichment analysis was performed by using the Functional Analysis module (MS peaks to pathways) in MetaboAnalyst v5.0. #SHE: number of significant hits enriched; #DCF: number of significant differential chemical features obtained by multiple unpaired t-test comparisons with Welch correction and Holm-Sidak post-hoc correction for multiple comparisons, setting an alpha threshold for significance at 0.01.

**Supplementary Dataset S5 (.xlsx):** Detailed output of pathway enrichment analysis performed by using the Functional Analysis module (MS peaks to pathways) in MetaboAnalyst v5.0.

**Supplementary Figure 4**. Pan-metabolome of cassava’s root tissue. **A**, score plot of components 1 and 2 of principal component analysis of root’s untargeted analysis, including over 500 chemical features detected as variables and 190 accessions as observations. **B**, scatter plot of chemical features’ relative abundance coloured by root metabotype (RM) clade. The most abundant features in each clade are annotated after putative identification of mass signals.

**Supplementary Dataset S6 (.xlsx):** putative identification of the most abundant chemical features per root metabotype clade.

**Supplementary Figure 5:** Metabolome stability of cassava roots over multiple crop years. **A**, Score plot of principal components 1 and 2 analysis of untargeted metabolite profiling of cassava roots accessions harvested from 2016 to 2020. Legend: green, 2016 crop; blue, 2017 crop; sky blue, 2018 crop; pale blue, 2019 crop; and red dots, 2020 crop. **B**, Orthogonal partial least square discriminant analysis (OPLS-DA) score plot comparing accessions harvested in 2016 (green dots) and 2020 (blue boxes). Table summarising model descriptors is presented at the top of the plot.

**Supplementary Figure 6:** Matrix correlation (Pearson) between whitefly phenotype and metabolites grouped by chemical class (right) and corresponding p-values (left).

**Supplementary Dataset S7 (.xlsx):** correlation (Pearson) and significance values between leaf chemical features and phenotypic records.

**Supplementary Dataset S8 (.xlsx):** correlation (Pearson) and significance values between root chemical features and phenotypic records.

