## Supplementary figures and images for "Integrated genetic and metabolic characterisation of diverse Latin American cassava (*Manihot esculenta* Crantz) germplasm; implications for future breeding strategies"

### Supplementary Figure 1

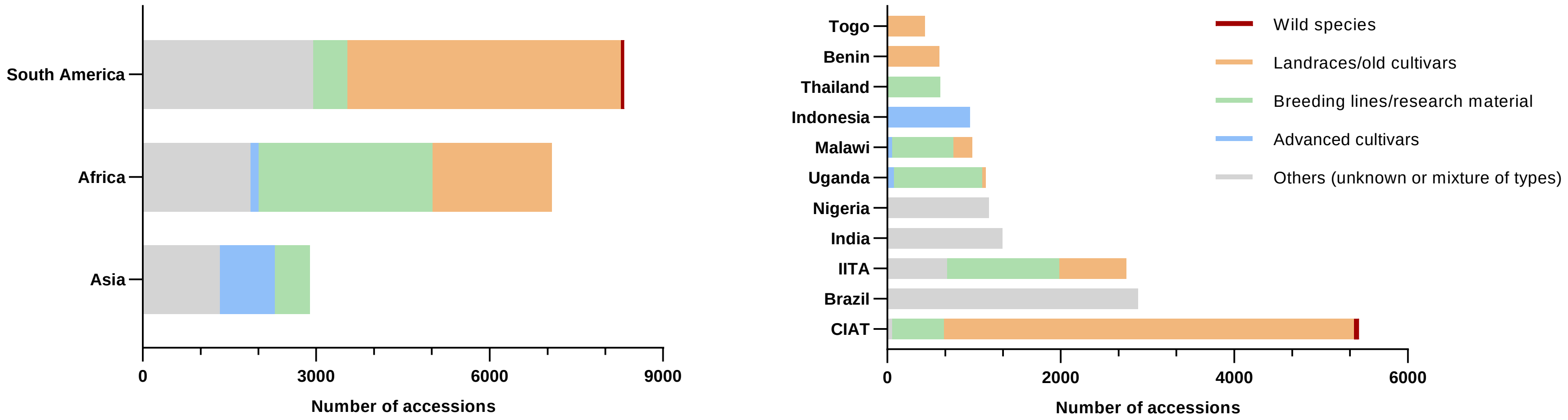

### Supplementary Figure 2

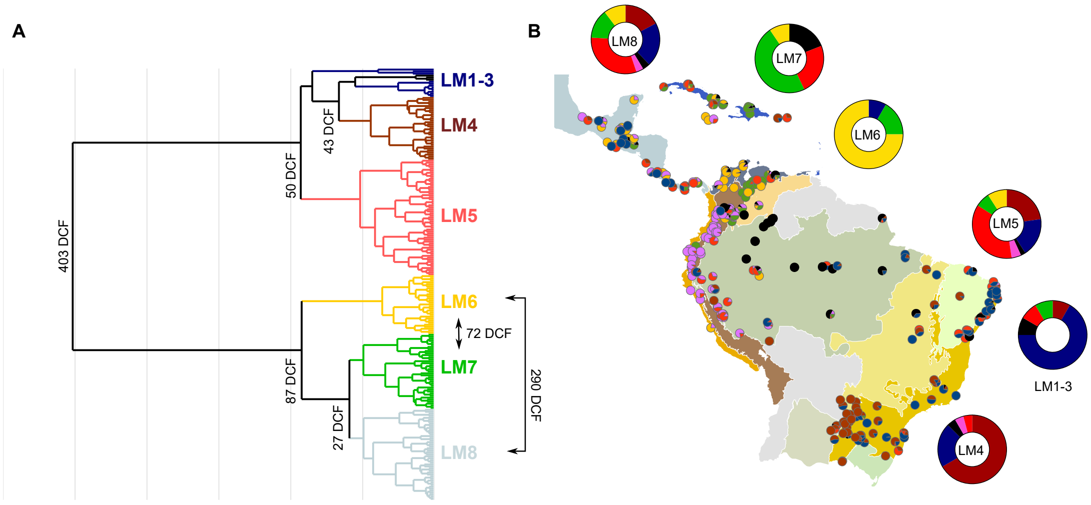

### Supplementary Figure 3

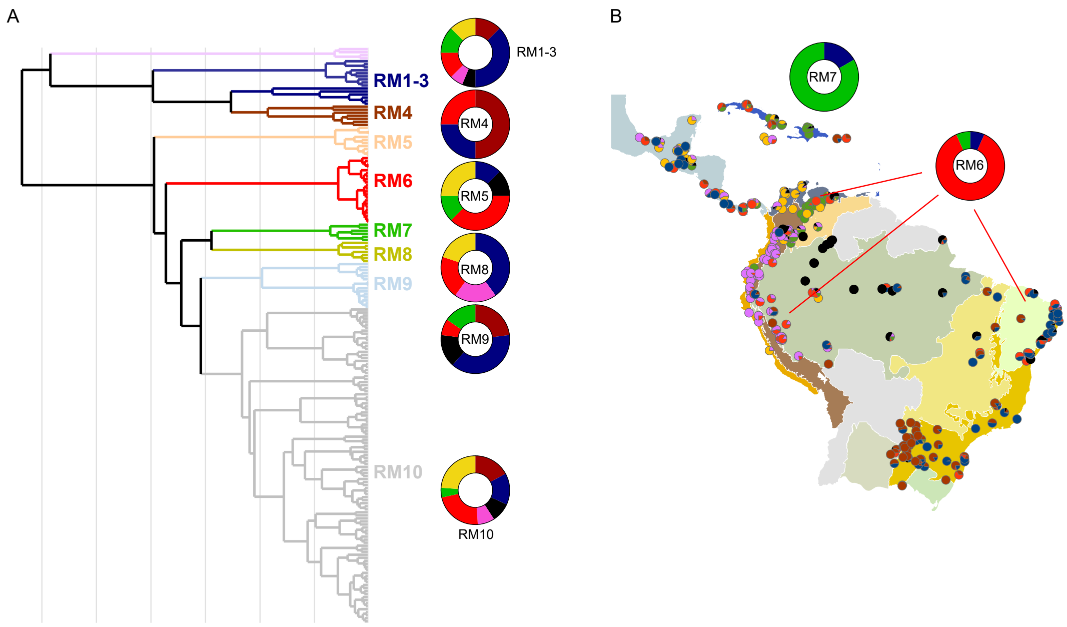

### Supplementary Figure 4

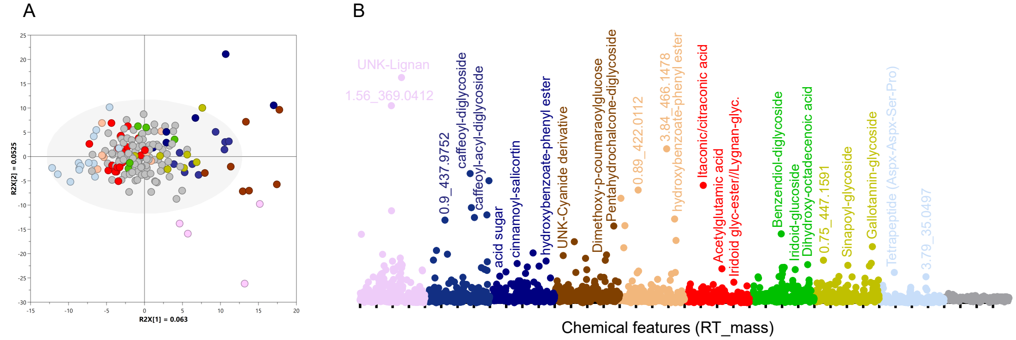

### Supplementary Figure 5

**A**

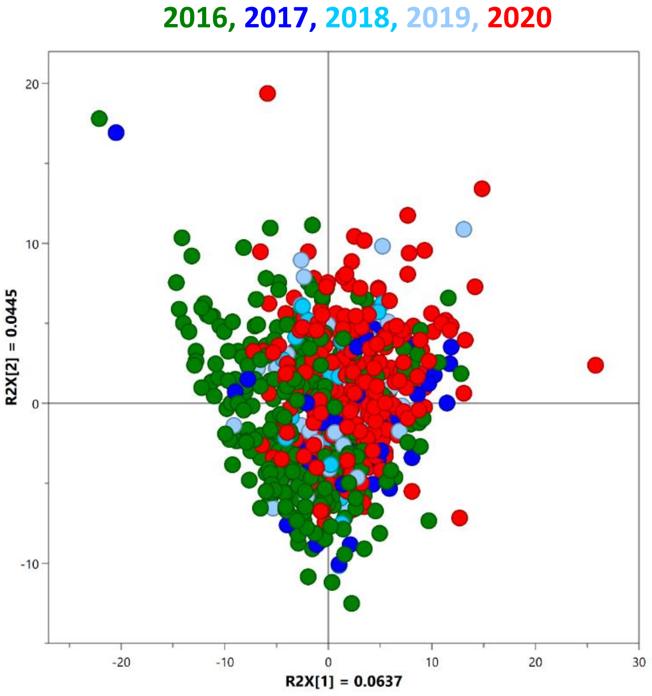


**B**

| R2X (X-axis): between class | R2X0 (Y-axis): within class | R2 (cum) | Q2 (cum) |
|---|---|---|---|
| 0.0233 | 0.0443 | 0.609 | 0.469 |

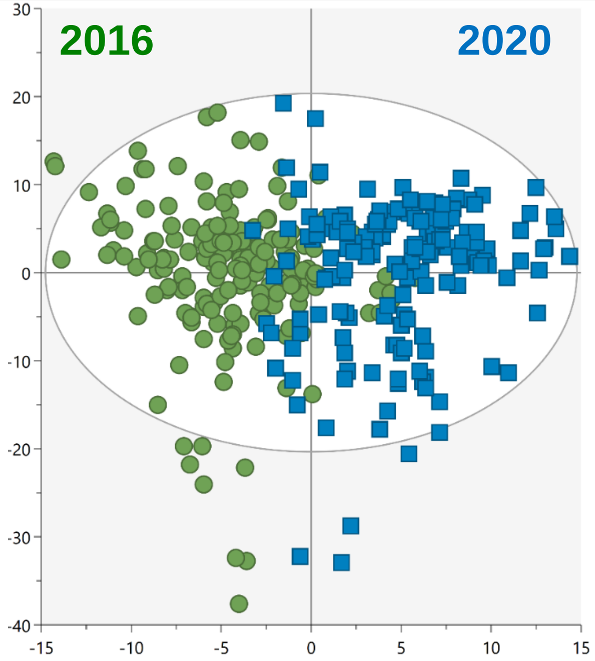

### Supplementary Figure 6

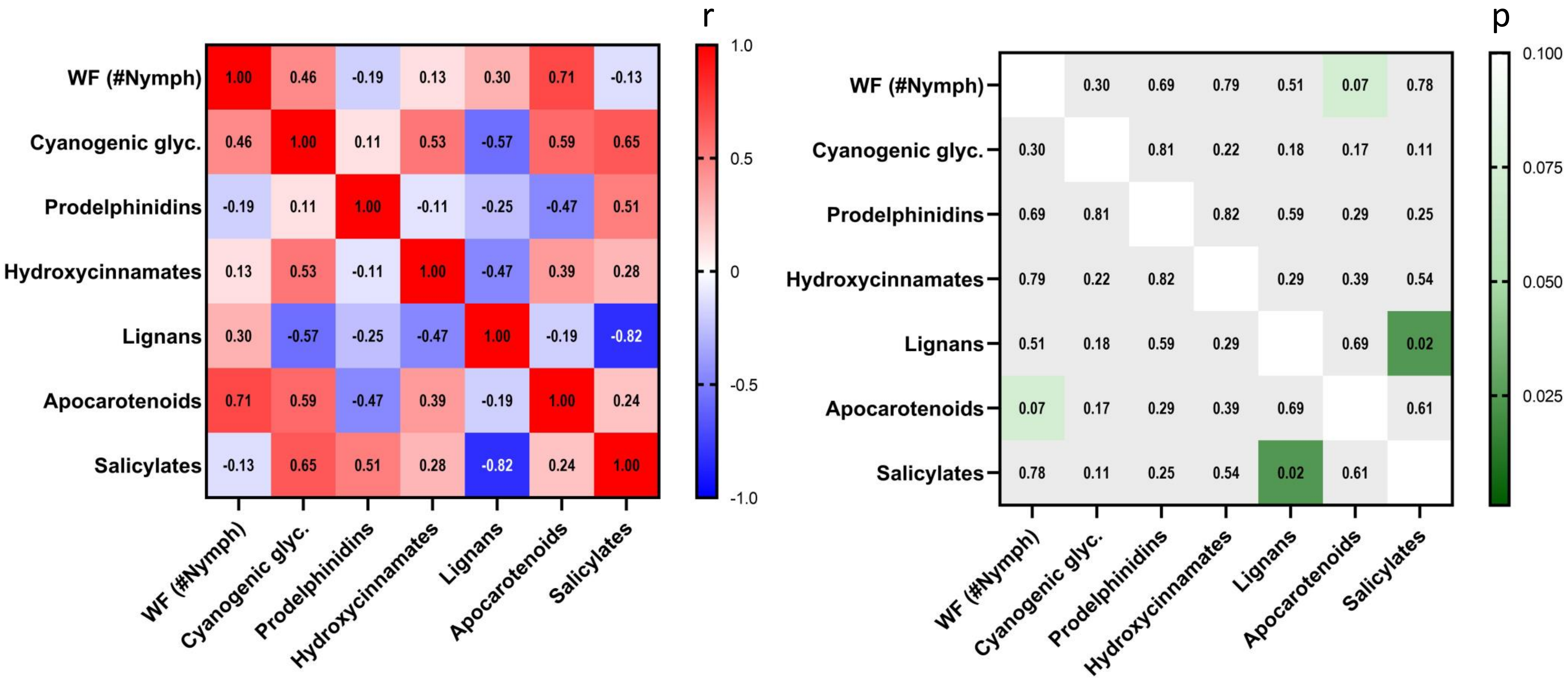
