## Supplementary Table 1 for "Integrated genetic and metabolic characterisation of diverse Latin American cassava (*Manihot esculenta* Crantz) germplasm; implications for future breeding strategies"

**Supplementary Table 1**: Summary of the most significant enriched pathways differentiating LM clades. Pathway enrichment analysis was performed by using the Functional Analysis module (MS peaks to pathways) in MetaboAnalyst v5.0. #SHE: number of significant hits enriched; #DCF: number of significant differential chemical features obtained by multiple unpaired t-test comparisons with Welch correction and Holm-Sidak post-hoc correction for multiple comparisons, setting an alpha threshold for significance at 0.01.

| **Pair-wise LM clades comparisons** | **# DCF** | **# SHE** | **Highest significant pathways enriched (ranked by permutation gamma p-value)** |
| --- | --- | --- | --- |
| 1. South (LM1-5) *Vs.* North (LM6-8) | 403 | 241 | - Flavonoid, flavone & flavonol, phenylpropanoids & anthocyanin biosynthesis - Val, Leu, Ile biosynthesis & degradation - Sugar metabolism |
| 2. Atlantic Forest (LM1-4) *Vs.* Mix-South (LM5) | 50 | 101 | - Riboflavin biosynthesis - Synthesis and degradation of ketone bodies - Lignans biosynthesis |
| 3. Dry (LM4) *Vs* Humid (LM1-3) Atlantic Forest | 43 | 54 | - Val, Leu, Ile biosynthesis - Organic acids |
| 4. Meso America (LM6) *Vs*. Savanna & Mix-North (LM7-8) | 87 | 53 | - Phenylalanine biosynthesis - Chlorophyll ring biosynthesis - Flavonoids, phenylpropanoids & anthocyanin biosynthesis |
| 5. Meso America (LM6) *Vs*. Savanna (LM7) | 72 | 64 | - Quinones biosynthesis - Lignan biosynthesis - Phe, Trp, Tyr biosynthesis |
| 6. Meso America (LM6) *Vs*. Mix-North (LM8) | 70 | 32 | - Chlorophyll ring biosynthesis - Flavonoids and phenylpropanoid biosynthesis |
| 7. Mix-South (LM5) *Vs*. Mix-North (LM8) | 290 | 129 | - Flavonoids and phenylpropanoid biosynthesis - Chlorophyll ring biosynthesis |
| 8. Savanna (LM7) *Vs*. Mix-North (LM8) | 27 | 33 | - Chlorophyll ring biosynthesis - Anthocyanin biosynthesis |
